# Perfringolysin O pore-forming complexes are predominantly integral multiples of six subunits

**DOI:** 10.1101/2024.06.20.598590

**Authors:** Meijun Liu, Xintao Qin, Menglin Luo, Yi Shen, Jiabin Wang, Jielin Sun, Daniel M. Czajkowsky, Zhifeng Shao

**Author notes:** Corresponding author. DM Czajkowsky, State Key Laboratory of Systems Medicine for Cancer, School of Biomedical Engineering, Shanghai Jiao Tong University, Shanghai, China.

## Abstract

Perfringolysin O is a well-studied bacterial cytolysin that forms large oligomeric pores with a wide range of sizes on membranes via a prepore intermediate. Here we examined the sizes of both PFO prepore and pore complexes electrophoretically using a multi-stack-gradient gel and found that, unexpectedly, there are only at most seven predominant sizes of either pores or prepores. Complexes extracted from each band exhibit contour lengths that are integral multiples of six subunits. High-resolution atomic force microscopy images of PFO pore complexes in supported bilayers also reveal a predominant hexameric-based stoichiometry. Thus, these results reveal a previously unknown structural hierarchy in PFO complexes, with larger complexes apparently built up from hexameric sub-complexes. We suggest that different inter-subunit interactions within and between the hexamers result in a likewise difference in the coordination of the prepore-to-pore transition within and between the hexamers, and is thus a critical feature of the allostery of this large multi-subunit complex.

## Introduction

Cholesterol-dependent cytolysins (CDCs) are a large family of bacterial pore-forming proteins (PFPs) that play significant roles in virulence^1–7^. These PFPs bind to the bilayer as monomers, self-assemble into prepore intermediates on the bilayer surface, and then cooperatively insert into the bilayer to form membrane-spanning pores complexes^8,9^. However, like other PFPs, it is not clear what triggers the transition from the prepore to the pore conformation and how activation of this trigger is coordinated among all of the subunits within such large complexes (apparently up to 50 subunits)^10–13^. Some other large biological complexes have been proposed to function via a nested hierarchical allostery^14^, exhibiting coordination within sub-assemblies at one level and then between sub-assemblies at a higher level, but the generality of this mechanism is still unknown^15^.

As for the trigger within the CDCs, an early suggestion was that pore formation occurred upon the completion of the full prepore ring^16^. Consistent with this, sodium dodecyl sulfate-agarose gel electrophoresis (SDS-AGE) of CDC complexes revealed only a single predominant oligomeric complex^16–21^. However, images of pore complexes obtained by atomic force microscopy (AFM), electron tomography, or electron microscopy have clearly shown heterogeneously sized arc-shaped complexes that are prominently present among complete rings^22–27^. Clearly, such a wide range of sizes would be expected to produce many bands in electrophoresis gels. Recent single-molecule fluorescence microscopy data also suggested that pore complexes can be incomplete rings^28,29^. Interestingly, this recent work also revealed that some complexes appear to continue to grow after pore formation, in apparent conflict with the aforementioned prepore model. Hence, resolution of this issue is expected to not only provide clarity into the identity of the bona fide CDC pore complex(es) but also reveal insight into the mechanism of pore formation.

Here, we re-investigated the sizes of the pore complexes of perfringolysin O (PFO), a well-studied member of the CDC family, by gel electrophoresis using a novel multi-stack polyacrylamide gel and indeed observed a range of sizes for both the prepore and pore complexes. However, unexpectedly, rather than a large number of bands or a smear, there were generally only at most 7 bands. AFM images of complexes isolated from these bands revealed that their contour lengths are integral multiples of 6 subunits. Moreover, AFM images of PFO pore complexes in supported lipid bilayers also revealed a predominant hexameric-based stoichiometry. Thus, overall, these results point to an unexpected hierarchy in the structure of PFO complexes, which recalls the nested hierarchical architectures of other large complexes. We propose that different inter-subunit interactions within a hexamer and between the hexamers result in a difference in the coordination of the prepore-to-pore transition within and between the hexamers.

## Results and Discussion

To reconcile the previous electrophoretic and imaging-based data, we sought to examine PFO complexes with a continuous gradient gel that we expected would better separate such large, differently-sized complexes than SDS-AGE. Pore complexes were examined by incubating PFO with cholesterol-containing lipid vesicles, followed by solubilization with gel-loading buffer (that contains 0.5% lithium dodecyl sulfate (LDS)) and then loading this into a 3-12% gradient polyacrylamide gel. However, as shown in Figure 1A, we also observed only a single dominant band of oligomers, similar to previous results obtained with SDS-AGE.

**Figure 1.**
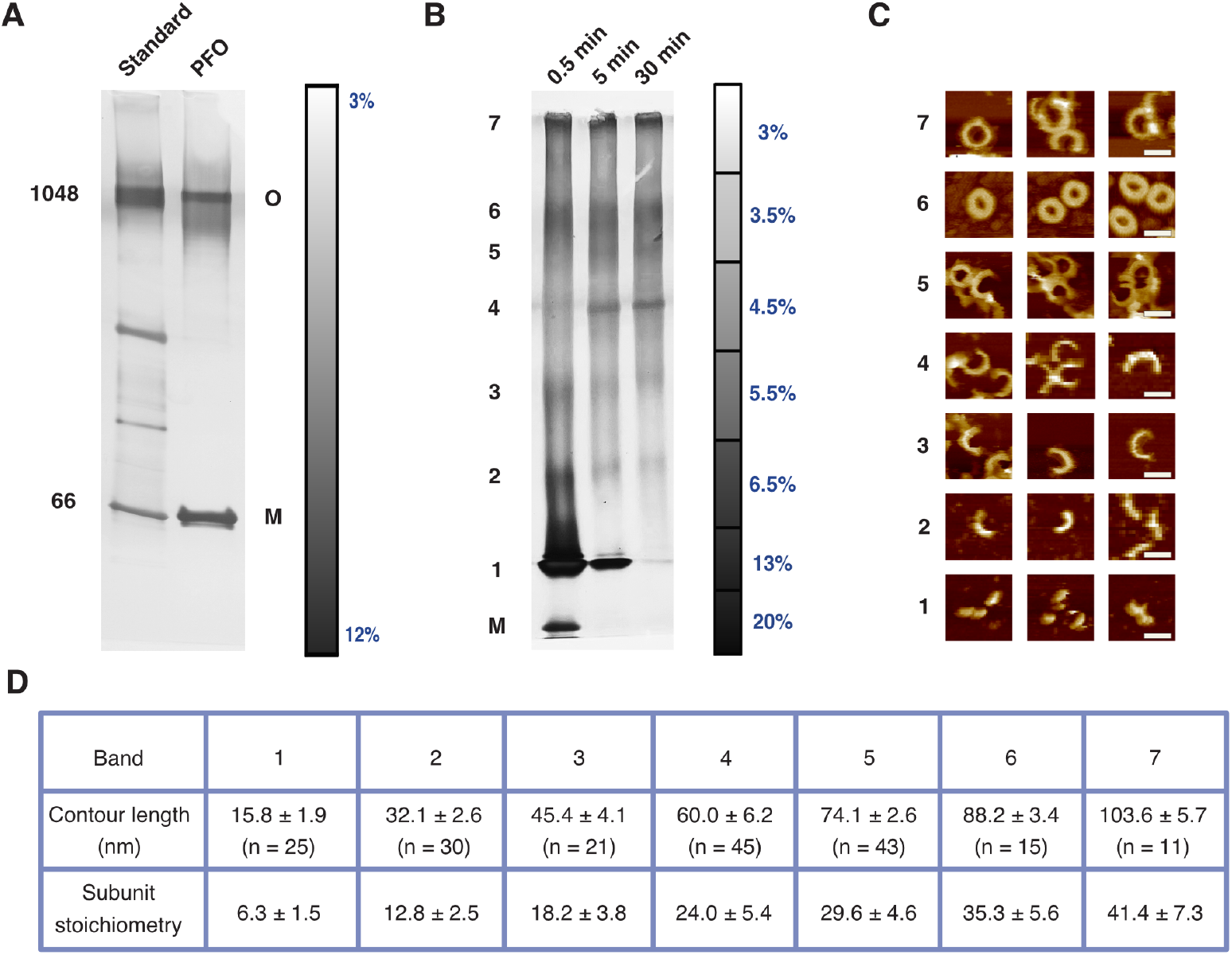
Gradient gel electrophoresis of PFO pore complexes. (A) A continuous 3-12% polyacrylamide gradient gel showing a predominant oligomeric band of PFO pore complexes in addition to the monomeric band (near 66 kDa). The scale bar reflects the gel density and the numbers indicate molecular weights of the standard (left lane). The “O” and “M” labeled on the right side of the gel refer to oligomer and monomer, respectively. (B) The multi-stack-gradient gel reveals maximally seven distinct bands of pore complexes. The relative dominance of the bands was found to change with PFO/vesicle incubation time. The density of the stacks in the scale bar is determined by the volume of each stack that was cast. (C) Typical AFM images of the complexes isolated from each of the individual bands (as numbered on the left in (B)). The complexes from the smaller five bands were predominantly incomplete rings, while the complexes in the larger two bands were mostly rings, with a minor portion of incomplete rings. Scale bar: 30 nm. (D) Measurements of the contour lengths from the individual bands. The lower row of subunit stoichiometry was obtained by dividing the contour length by the circumferential length of a single subunit (2.5 nm).

We hypothesized that the resolving power of this gel might still not be sufficiently high enough to separate the differently sized complexes and thus systematically examined a number of gel-casting conditions (Supplementary Figure S1). Ultimately, we found that sequential stacks of different polyacrylamide density yielded the clearest separation of multiple well-defined bands (Figure 1B, Supplementary Figure S1, Methods). However, instead of a multitude of bands, these gels generally showed only a very few dominant bands. We found that the presence of some bands or their relative dominance changed with protein/vesicle incubation time (Figure 1B, Supplementary Figure S2), gel density (Supplementary Figure S1), and other sample conditions (Supplementary Figure S2). However, the maximum number of bands that we routinely resolved was seven (Figure 1B, Supplementary Figure S2).

To determine the physical properties of the complexes in these bands, we extracted the complexes from each gel-resolved band, deposited them on mica, and imaged them with AFM in solution. As shown in Figure 1C, overall, these images revealed both incomplete- and complete-ring complexes, with those complexes from the lower-weight bands exclusively arc-shaped and the higher-weight bands consisting mostly of complete rings. Thus, consistent with previous imaging-based observations^22–29^, gel electrophoresis indeed can resolve the PFO pore complexes as a range of differently sized complexes that are either incomplete- or complete-rings. These AFM images were not of a sufficient resolution to resolve individual subunits within the complexes, likely owing to a poorer resolving capacity of AFM at sparse surface coverages of proteins on the substrate^30^. Thus, we estimated the stoichiometry of these complexes based on their contour lengths using the known circumferential subunit length of the pore complexes (2.5 nm^23,31^). We found that the contour lengths within each band were narrowly defined, with sequential bands consisting predominantly of ∼6, 12, 18, 24, 30, 36 and 42 subunits, respectively (Figure 1D). That is, overall, the PFO pore complexes are predominantly integral multiples of 6 subunits.

To verify this observation, we sought to directly image PFO pore complexes within supported lipid bilayers by AFM^32–35^. As previously observed^23^, images of such complexes revealed both incomplete-ring and complete-ring complexes of a range of different sizes and curvatures with sufficiently high resolution to resolve individual subunits (Figure 2A). Directly counting the stoichiometry from such images indeed revealed a predominance of complexes that are integral multiples of 6 subunits (Figure 3).

**Figure 2.**
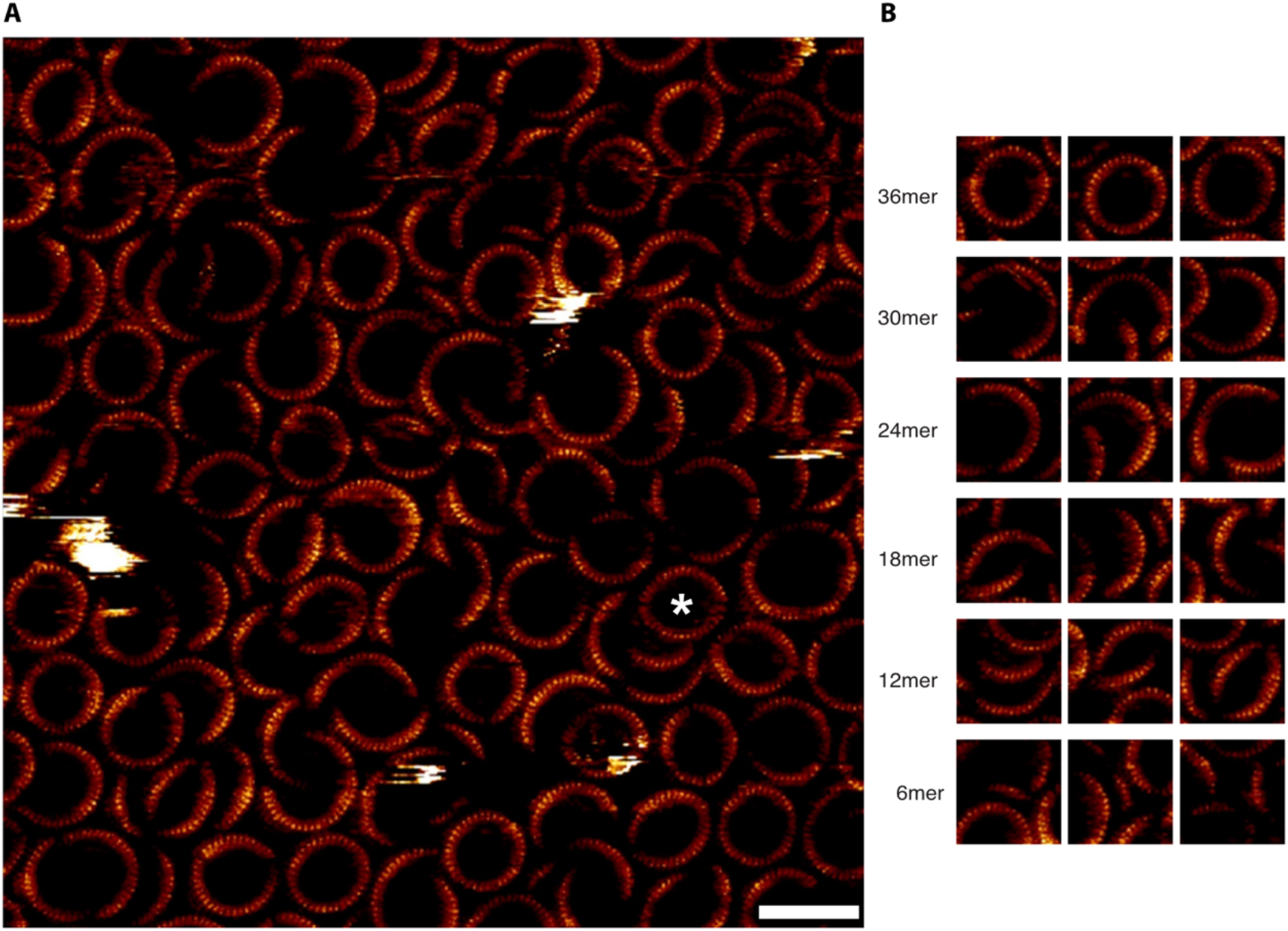
High resolution AFM images of PFO pore complexes in supported lipid bilayers directly resolves subunit stoichiometry. (A) Large scale image of the complexes in the supported bilayer. The asterisk indicates an example of a complex that appears to be formed by the apposition of two arcs of a similar curvature as in previous studies^16,22–27^. Scale bar: 40 nm. (B) Isolated images showing the subunit stoichiometry indicated on the left.

**Figure 3.**
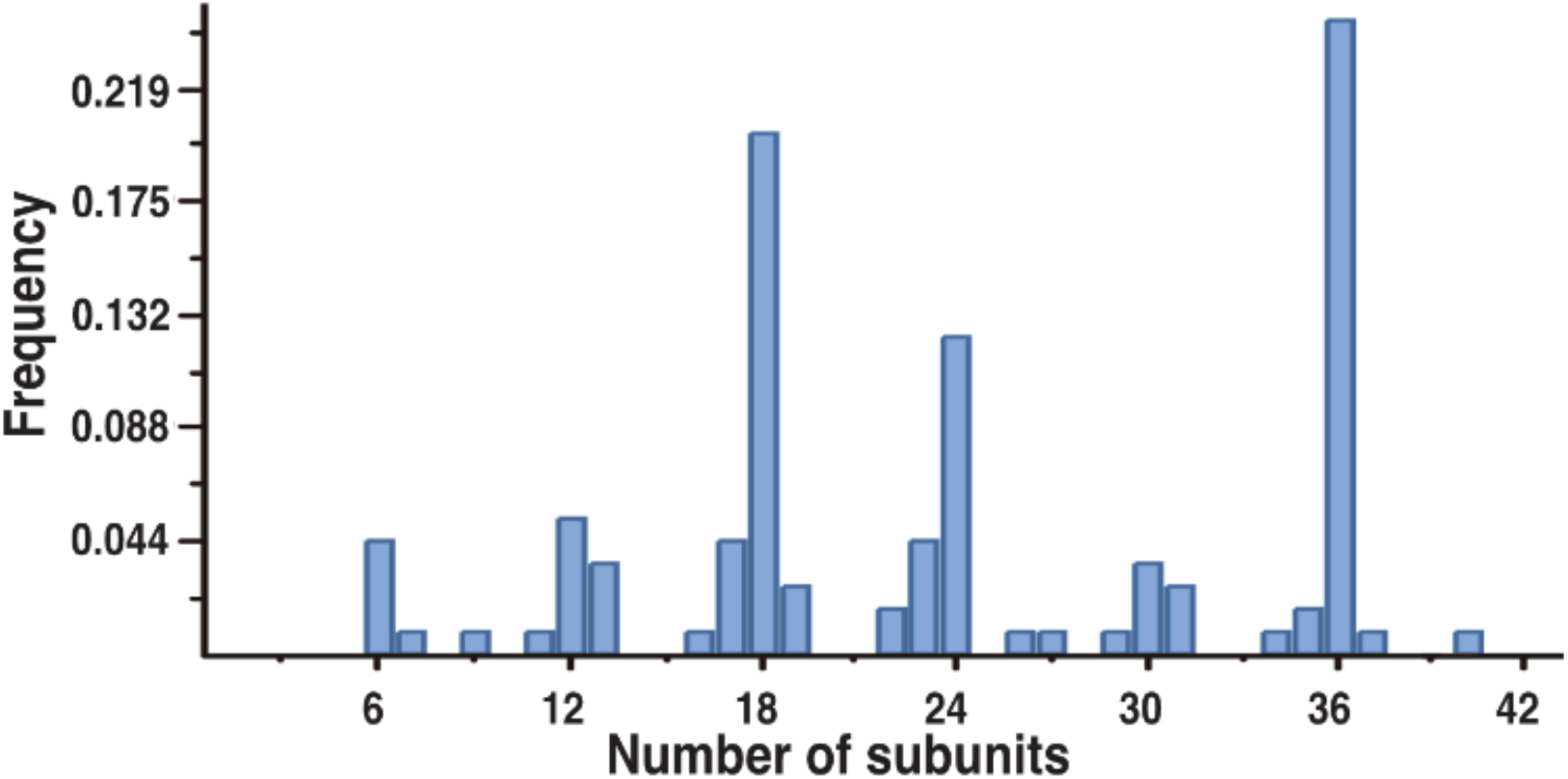
Overall distribution of subunit stoichiometry of PFO pore complexes determined from high-resolution AFM images.

We note that the aforementioned initial SDS-AGE study^16^ also provided single-channel electrophysiological data that appeared to indicate a single predominant conductance of the complexes. Thus, we re-examined this experiment as well by studying single channels formed directly in cholesterol-containing planar lipid bilayers (Methods). However, rather than a single predominant conductance, we observed a wide-range of conductances (Supplementary Figure S3), consistent with the formation of a wide-range of pore complex sizes observed with AFM and in the multi-stack gels. Thus, altogether, these results show that PFO pore complexes are predominantly integral multiples of 6 subunits. To determine whether this feature is present even at the prepore stage, we examined the sizes of prepore-trapped intermediates with the multi-stack gel. As Figure 4A shows, prepore-trapped complexes formed via either of two different ways (namely, low temperature with wild-type protein or using disulfide-trapped mutants) exhibit the same few bands as the pore complexes, also maximally seven. Thus, these results indicate that there is a striking preference for hexameric-based complexes in both the prepore and pore states.

**Figure 4.**
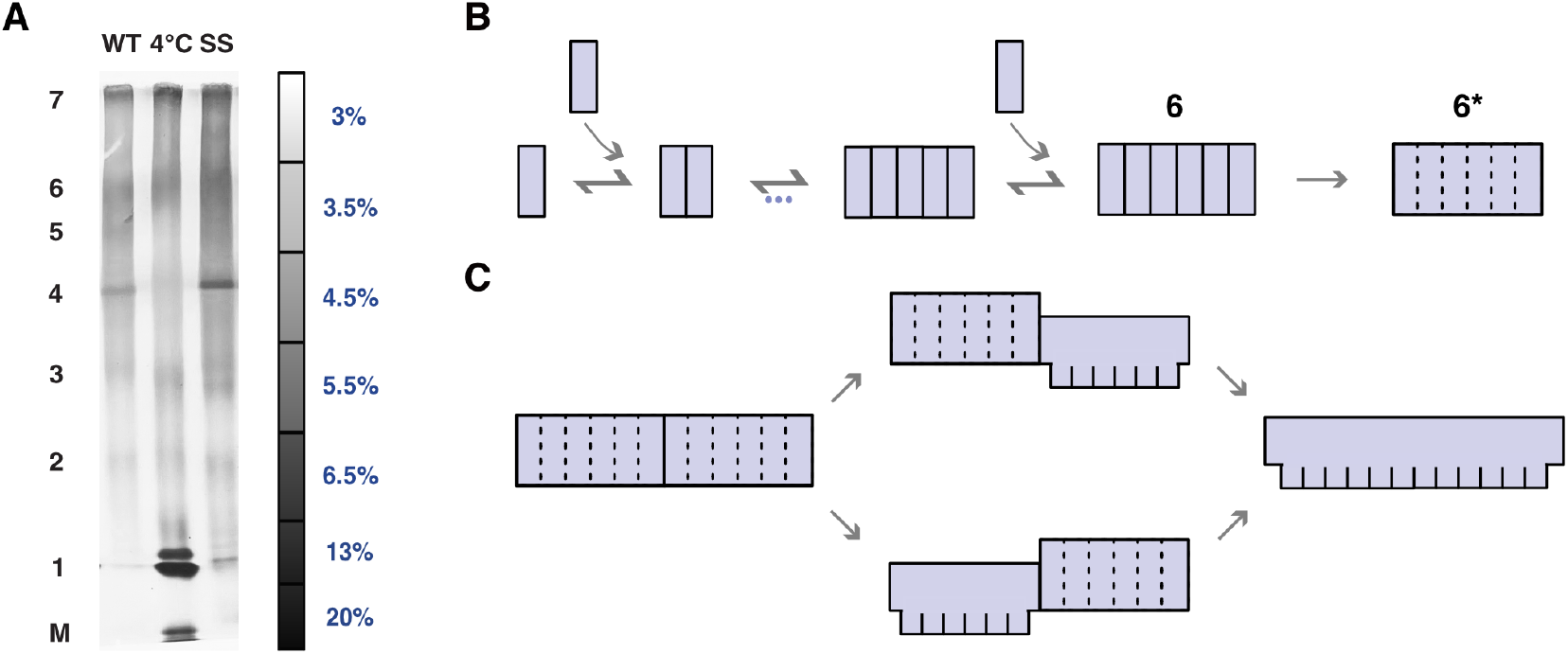
Multi-stack-gradient gel of prepore complexes provides support for a novel model of the pore-forming mechanism. (A) Prepore complexes (PFO^G57C/S190C^, labeled SS in the figure^42^, and wild-type PFO at lower temperature (4°C)^16^) resolve into the same few bands (maximally seven) that were observed with the pore complexes. The wild-type PFO lane to the left is the same lane to the right in Figure 1B to enable comparison. (B) Schematic model to explain observations presented here, whereby upon forming hexamers, the complexes undergo a transition to a structure in which subunits now exhibit highly cooperative behavior (depicted by the dotted lines). (C) Proposed model in which the structural transition from prepore-to-pore occurs cooperatively within each hexamer, with a weaker coordination between neighboring hexamers (separated by solid line).

Hence, overall, these results reveal a hierarchical subunit organization within both PFO prepore and pore complexes. Further, it suggests the presence of an important step upon hexamerization within the prepore state that persists to the pore state. In particular, we propose that monomers assemble on the membrane until forming hexamers, at which point they undergo a transition to a more cooperative (detergent-resistant^36^) hexameric prepore complex (Figure 4B). In our model, conversion to the pore conformation would then occur simultaneously within this hexamer, with neighboring hexamers transitioning either independently or with reduced cooperativity than within the hexamers (Figure 4C). As such, we suggest that PFO complexes undergo the prepore-to-pore transition by a nested model of allostery^14^, with stronger coordination within the hexamers than between them. With this, during pore formation, any single complex (larger than a hexamer) would be at least transiently composed of sub-complexes in the prepore state as well as those in the pore state (simultaneously), which has indeed been recently observed with high-speed AFM imaging of another CDC member^37^ and in our earlier study of a disulfide-trapped PFO mutant (Supplementary Figure S4).

Finally, we note that there are two possible mechanisms by which complexes grow in size after attaining this cooperative hexameric prepore state: with monomers continuing to bind to this prepore or with exclusive interactions between cooperative hexamers (Supplementary Figure S5). Of note, the former mechanism affords for the possibility of monomers binding to a complex that has already partially transitioned to the pore conformation, which could explain the continued growth of a pore complex observed in recent single-molecule fluorescence microscopy studies^28,29^.

## Conclusions

Based on the analysis of both gel electrophoretic data and high-resolution AFM images, we have shown, for the first time, that PFO pore complexes are predominantly multiples of 6 subunits, which we anticipate plays a critical role in coordinating structural changes between the subunits to the pore state. Thus, unlike what is commonly believed, the subunits within a complex are not all equivalent, with those within a hexamer interacting in a different way than those between hexamers. At the same time, these results re-focus attention to a major unresolved issue that the only-complete-rings model was purported to resolve: the trigger to pore formation, which indeed remains unanswered for many PFPs. In our model, this trigger occurs following hexamerization and predominantly at the level of this hexamer, but the nature of this trigger indeed remains unresolved. We anticipate that future studies of this hexamer should provide mechanistic insight to this critical question. Further, we anticipate that resolution of this problem will also shed light into the more general question of structural coordination mechanisms within nested protein architectures that underlie the functioning of higher-order biological assemblies more broadly.

## Materials and Methods

### Expression and purification of PFO and its derivative

The sequences for the N-terminal His-tagged wild-type PFO and the disulfide-locked mutant PFO^G57C/S190C^ were inserted into the pPROEXHTα vector by Sangon Biotech. The *E.coli* BL21(DE3) cells (TransGen Biotech) containing recombinant plasmid were grown in LB medium with 0.01% ampicillin at 37°C overnight, transferred into LB medium (1:100), grown until attaining an OD value of 0.8, and then induced with 1 mM isopropyl β-D-1-thiogalactopyranoside (IPTG, Sangon Biotech) at 16°C for 18 hours. Cells were harvested at 6000 rpm for 5 minutes, resuspended in PBS and frozen-thawed for 3 cycles, after which the lysis buffer (0.15ml CelLytic B (Sigma-Aldrich), 0.0015g Lysozyme, 15 μl PMSF and 0.5 μl Benzonase Nuclease (Yeasen) in 1.33 ml PBS) per 1 g wet cell pellet was added. The supernatant was incubated with ProteinIso Ni-NTA agarose Resin (TransGen Biotech) (2000:1), rotating at 4°C for 2 hours. The resin was then washed thoroughly with 20 mM imidazole in PBS and then eluted with 300 mM imidazole in PBS. The protein solution was dialyzed and stored with 20% glycerol at -80°C. The purity was evaluated with SDS-PAGE and hemolytic activity. For the latter, in short, PFO was incubated with about 1 × 10^9^ erythrocytes from washed rabbit blood at 37°C for 30 min. The percentage of hemolysis was determined by the release of hemoglobin monitored by the absorbance at 540 nm. The amount of toxin that produced 50% hemolysis was similar to published values^31^.

### Vesicle preparation

Cholesterol, egg L-α-phosphatidylcholine (eggPC) and heart total lipid extract were purchased from Avanti Polar Lipids. The lipid powder was dissolved in chloroform to a stock concentration of 25 mg/ml. The solvent mixture was evaporated using nitrogen gas to form a homogenous dry lipid film. The lipid film was resuspended in 150 mM NaCl, 20 mM Hepes (pH 7.4) to a final concentration of 1 mg/ml and then sonicated to form unilamellar vesicles.

### Native gradient PAGE

We first examined PFO pore complexes using continuous gradient polyacrylamide gel electrophoresis (PAGE) for finer separation of differently sized complexes than agarose^38,39^. The 3-12% native gradient gel in Figure 1A was made using an automatic gradient former and the NativePAGE 3-12% Bis-Tris gel in Supplementary Figure S1A was purchased from Invitrogen. PFO and the vesicle solution containing eggPC:cholesterol 1:1 (molar ratio) were incubated at 37°C for 30 minutes followed by the addition of the NuPAGE LDS 4× sample buffer (Invitrogen). The solubilized complexes and the NativeMark unstained protein standard (Invitrogen) were then loaded into the gel. Running conditions were 150 V for 60 min and 250 V for 30 min in a XCell SureLock Mini Cell (Invitrogen) using NuPAGE Tris–Acetate SDS running buffer, pH 8.25 (Invitrogen). Gels were stained by silver stain.

### Multi-stack gel electrophoresis

The gel was cast manually, stack by stack, with a minimum difference in cast gel density of 0.5% and a volume of 0.4 ml between neighboring stacks. A maximum number of 10 gel stacks could be cast for a 1.0 mm thick plate, and, overall, we examined stacks with gel densities between 3% to 20%. The reagents used were free of reducing agents, and the samples were not boiled. A stock polyacrylamide solution of 40%T (T, weight percentage of the acrylamide) and 4%C (C, weight percentage of crosslinker) was pre-cooled and APS and TEMED were both added to, at most, 0.1% (v/v). The casting process was finished in 2 minutes using a wide pipette tip. The sample preparation procedure of PFO and its derivative was the same as described above. A tris-tricine running buffer system (50 mM Tris, 50 mM Tricine, pH 8.3) was used. The gels were run by applying 26 V for 30 minutes, 50 V for 2h, and then 70 V for 10 h to maintain a maximum current of 0.1 A. Gel staining was the same as described above for the native gradient PAGE.

### Gel isolation and imaging

Since the proximity of folded proteins to molecular standards in gels does not provide a reliable measure of their size, we determined the size of the complexes by first isolating the complexes from each band and then imaging with AFM^40^. For this, one lane of the gel containing the sample was removed and stained to identify the bands. This lane was then put back into the gel and used as a marker to identify the appropriate positions within a lane of sample that was not stained. Each plug containing a single band was then placed into 2 ml centrifuge tubes and was ground manually and then placed in 1.5 ml water. The sample was then shaken for two days at 9°C, and then concentrated by a 100 kDa ultrafilter prior to AFM imaging. For AFM, the sample was applied to freshly-cleaved mica and incubated for 30 minutes, followed by washing thoroughly with 150 mM NaCl, 20 mM Hepes, 40 mM CaCl2 (pH 7.4). The contour lengths were measured at the centers of the arc/ring complexes. To account for the effect of tip broadening on this measurement, we estimated the magnitude of the tip broadening effect based on the (radial) topographical profile within the middle of the complexes. We then subtracted this distance from each end of the contour to obtain the final contour lengths. AFM imaging was performed using a Multimode 8 AFM (Bruker) at room temperature in the contact mode with the “E” scanner with 0.06 N/m cantilevers (SNL-10 probe, Bruker). Images were recorded at a scan rate of 6 Hz and analyzed by Nanoscope Analysis (Bruker).

### AFM sample preparation and imaging

The bilayer was made by forming two monolayers on mica using Teflon well as described in previous study^41^. The first monolayer consisted of egg PC, and the second monolayer was eggPC:cholesterol 1:1 (molar ratio). The bilayers were rinsed with imaging buffer consisting of 150 mM NaCl, 20 mM Hepes, 40 mM CaCl2 (pH 7.4) before 1 μg PFO was injected into the subtending solution, with a final protein concentration of 20 μg/ml. Following incubation, the excessive monomers were washed away thoroughly before imaging. With PFO^G57C/S190C^, 5mM DTT was added into the solution before imaging for Supplementary Figure S4. AFM imaging was performed as described above.

### Planar Bilayer Membrane Measurements

All electrophysiological measurements were performed with an BC-535 bilayer clamp amplifier (Warner Instruments) paired with an LPF-8 8-pole Bessel Filter (Warner Instruments). Planar membranes were made from decane solutions of 1-palmitoyl-2-oleoyl-*sn*-glycero-3-phosphocholine (POPC) and cholesterol mixed at a ratio of 45:55 mol%, and total lipid concentration was 20 mg/mL. Membranes were formed across a 200 μm diameter hole in a Teflon cup, separating the apparatus into two chambers. Each chamber was filled with a 1 mL buffer (0.1 M NaCl, 10 mM HEPES, pH 7.0) and a pair of Ag/AgCl electrodes were used to record transmembrane current with a holding potential of 50 mV. Afterwards, 6.5 μg PFO monomers incubated with 0.2% DDM were added to the cis chamber (1:1000 dilution) to form pores in the membrane. All channel signals were recorded with a low pass filter frequency of 5 kHz and a sampling frequency of 25 kHz. The data were analyzed with Clampfit10.4.

## Author Contributions

Meijun Liu performed most of the experiments and was a main author of the manuscript; Xintao Qin performed the electrophysiological measurements and analysis; Menglin Luo performed some of the AFM imaging; Yi Shen provided guidance for the AFM experiments and analysis; Jiabin Wang contributed to the protein purification; Jielin Sun helped supervise the project; Daniel M. Czajkowsky conceived the project, supervised the work, and was a main author of the manuscript; and Zhifeng Shao was a major supervisor of the work. All authors read and approved of the final version of the manuscript.

## Acknowledgments

This work was supported by grants from the National Key R&D Program of China (No. 2020YFA0908100), the National Natural Science Foundation of China (Nos. 31971151, 81627801, 81972909, and 32370572), and the K.C. Wong Education Foundation (H.K.). We thank Ming Cheng for his help with these experiments.

