## Supplemental figures for "Perfringolysin O pore-forming complexes are predominantly integral multiples of six subunits"

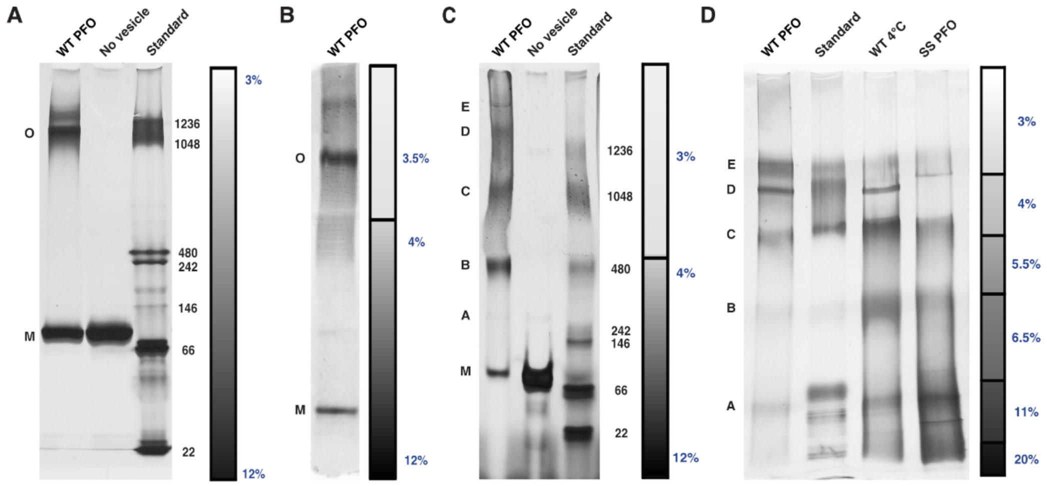


**Supplementary Figure S1.** Detection of different numbers of bands in gels of different compositions. (A) Only two bands, one associated with the PFO monomer and the other associated with PFO oligomers, are evident in a continuous gradient 3%-to-12% Bis-Tris NativePAGE gel (Invitrogen). (B-D) Changing the composition of the gels result in the resolution of more than a single band of the oligomers. The density of the stacks in the scale bar is determined as in Figure 1B. The dominant resolved bands are labelled by letters. Ultimately, the multi-stack-gradient gels with the appropriate densities (as labeled) yielded the greatest number of distinct bands.

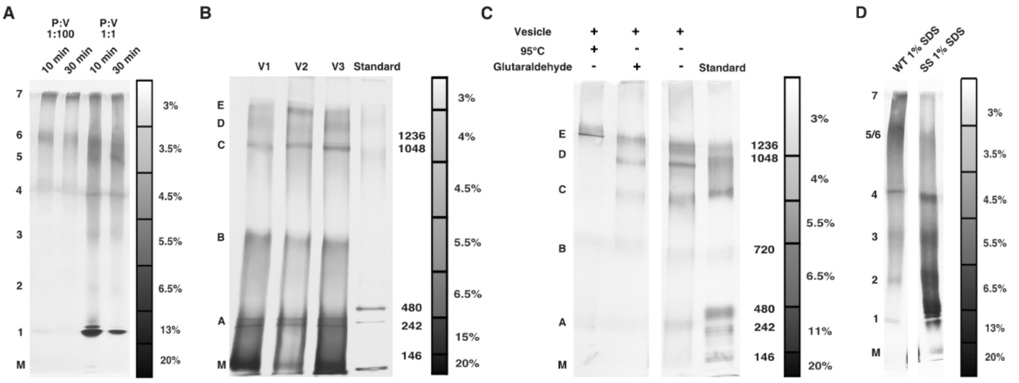


**Supplementary Figure S2.** Different sample conditions examined with the multi-stack gradient gels. (A) PFO pore complexes with 1:1 (lane 3-4) and 1:100 (lane 1-2) protein-to-lipid mass ratios with different protein/vesicle incubation times, PFO pore complexes with 1:10 protein-to-lipid ratio is shown in Figure 1B. (B) PFO pore complexes formed on vesicles with different lipid compositions: heart total extract:cholesterol 1:2 (molar ratio) (V1), eggPC:cholesterol 1:1 (molar ratio) (V2), and heart total extract:cholesterol 1:1 (molar ratio) (V3), respectively. (C) Additional sample conditions that were used in the earlier SDS-AGE study[16]. (D) The pore and prepore complexes treated with 1% SDS show the degree of dissociation caused by this detergent.

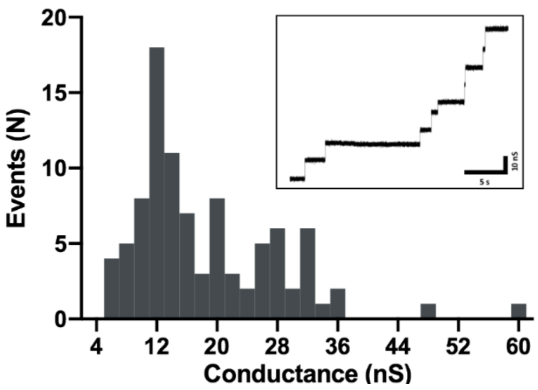


**Supplementary Figure S3.** Histogram of the distribution of channels formed by PFO in POPC-cholesterol planar bilayers from 11 separate planar membrane experiments (93 channels were recorded in total). From these measurements, 66% of the channels were examined to have a conductance between 10 to 20 nS (approximately 18 - 30 nm in diameter of the pore complexes). A typical example of the channels formed by PFO is shown in the inset of this figure.

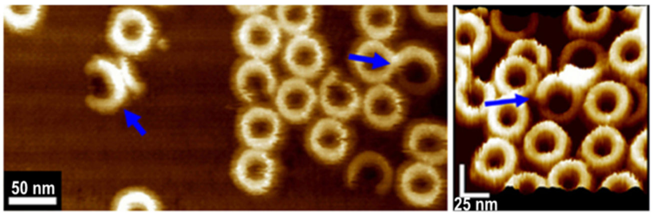


**Supplementary Figure S4.** AFM images of PFO$^{G57C/S190C}$ show the co-occurrence of prepore and pore sub-complexes within a common complex after a brief incubation with DTT, followed by fixation with 2% glutaraldehyde. The blue arrows point to complexes with both taller (prepore) and shorter (pore) sub-complexes simultaneously. Notice that the complex on the left in the left panel indeed appears to form a pore in the membrane, as evidenced by the absence of bilayer within the region delimited by this complex.

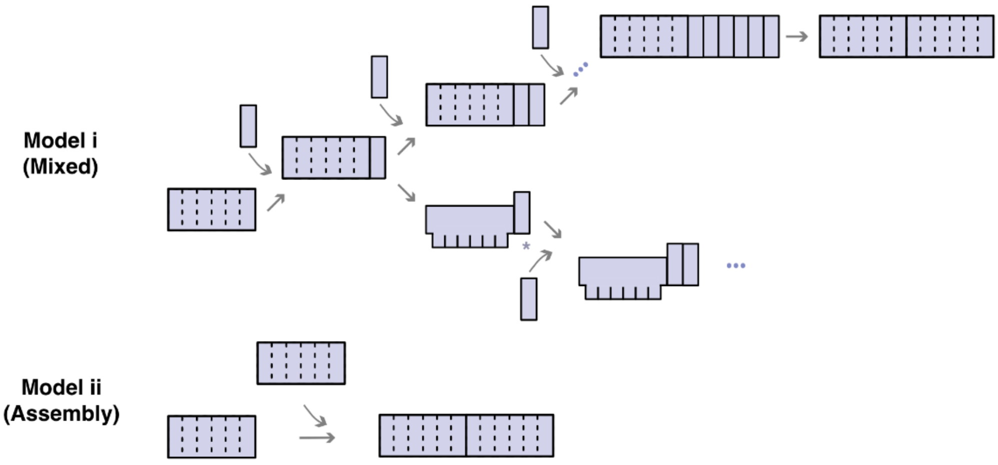


**Supplementary Figure S5.** Schematic diagram showing two mechanisms that reflect how complexes could possibly grow after achieving the cooperative hexameric prepore state (dotted line). Model i ("Mixed") reflects monomers continuing to bind to the hexameric prepore and Model ii ("Assembly") reflects formation of larger complexes exclusively via the interaction between cooperative hexamers. With the former, newly added monomers would continue to bind until attaining a hexameric stoichiometry, after which it would transition to a cooperative hexameric sub-complex. Prior to attaining this cooperative hexameric structure, there is a weaker degree of interaction between these monomers, which (in this model) would disassemble upon solubilization with detergent and thus complexes that are not hexameric are not observed in the multi-stack-gradient gels. In this scenario, it would also be possible for the first-assembled cooperative hexamer to transition to the pore conformation prior to formation of the second cooperative hexamer.
